# TET2 binding to enhancers facilitates transcription factor recruitment in hematopoietic cells

**DOI:** 10.1101/336008

**Authors:** Kasper D. Rasmussen, Ivan Berest, Sandra Kessler, Koutarou Nishimura, Lucía Simón-Carrasco, George S. Vassilou, Marianne T. Pedersen, Jesper Christensen, Judith B. Zaugg, Kristian Helin

**Affiliations:** Biotech Research and Innovation Centre (BRIC), Faculty of Health and Medical Sciences, University of Copenhagen, 2200 Copenhagen, Denmark.; The Novo Nordisk Foundation Center for Stem Cell Biology (Danstem), Faculty or Health and Medical Sciences, University of Copenhagen, 2200 Copenhagen, Denmark.; European Molecular Biology Institute, Structural and Computational Unit.; Wellcome Trust Sanger Institute, Wellcome Trust Genome Campus, Cambridge, United Kingdom and Department of Haematology, Cambridge University Hospitals NHS Trust, Cambridge, United Kingdom.; Friedrich Miescher Institute for Biomedical Research (FMI), CH-4058 Basel, Switzerland.; Centre for Gene Regulation and Expression (GRE), School of Life Sciences, University of Dundee, Dundee, UK.

**Keywords:** Acute Myeloid Leukemia, Chromatin, DNA binding, DNA methylation, Epigenetics, Hematopoiesis, TET2, Transcription Factor

## Abstract

The epigenetic regulator *TET2* is frequently mutated in hematological diseases. Mutations have been shown to arise in hematopoietic stem cells early in disease development, lead to altered DNA methylation landscapes and to an increased risk of hematopoietic malignancy. Here, we show by genome-wide mapping of TET2 binding sites in different cell types that TET2 localizes to regions of open chromatin and cell-type specific enhancers. We find that deletion of *Tet2* in native hematopoiesis as well as fully transformed Acute Myeloid Leukemia (AML) results in changes in transcription factor (TF) activity within these regions, and we demonstrate that loss of TET2 leads to enzymatic activity-dependent attenuation of chromatin binding of the hematopoietic TF CDX4. Together, these findings demonstrate that TET2 activity shapes the local chromatin environment at enhancers to facilitate TF binding and provide a compelling example of how epigenetic dysregulation can affect gene expression patterns and drive disease development.

## INTRODUCTION

The Ten-Eleven-translocation (TET) enzymes (TET1-3) mediate active DNA demethylation of cytosines in CG dinucleotides. This occurs by processive TET-mediated oxidation of 5-methylcytosine (5mC) to 5-hydroxymethylcytosine (5hmC), 5-formylcytosine (5fC) and 5-carboxycytosine (5caC). The presence of 5hmC may lead to passive replication-dependent loss of DNA methylation while 5fC and 5caC can be excised by thymine-DNA glycosylase (TDG) and replaced by unmodified cytosine via the base-excision repair (BER) pathway. While targeting of TET1 to chromatin has been investigated (Williams et al. 2011; Wu et al. 2011), the mechanisms of TET2 recruitment to chromatin remain poorly understood (reviewed in (Rasmussen and Helin 2016)).

Loss-of-function mutations of *TET2* have been found in patients with a wide range of hematological diseases, including AML (Scourzic et al. 2015). More recently, high frequencies of *TET2* mutations have also been observed in aging-associated Clonal hematopoiesis (CH) (Jaiswal et al. 2014; Genovese et al. 2014; Xie et al. 2014) and in the poorly studied disorder Clonal cytopenia of unknown significance (CCUS) (Kwok et al. 2015; Hansen et al. 2016). In previous studies, we and others identified a role of TET2 in protecting enhancer elements from aberrant DNA methylation (Lu et al. 2014; Hon et al. 2014; Rasmussen et al. 2015; An et al. 2015; Yamazaki et al. 2015). In addition, inhibition of TET proteins was shown to perturb chromatin architecture at enhancers in an embryonal carcinoma cell line undergoing neuronal differentiation (Mahé et al. 2017). Despite of these compelling results, direct TET2 binding at enhancers has not been reported. In fact, previous studies mapping TET2 genome-wide occupancy noted significant TET2 binding at CpG islands and promoters (Chen et al. 2013; Deplus et al. 2013; Peng et al. 2016). These seemingly contradictory observations as well as the impact of aberrant DNA methylation at enhancer elements in hematopoietic cells remains to be resolved.

Gene expression is regulated by transcription factors (TFs) that bind DNA in a sequence-specific manner. Activation of a specific gene locus depends both on concentration of individual TFs as well as their ability to access regulatory elements, often defined as enhancers, that control gene expression. TF binding outside gene promoters is associated with low- or intermediate DNA methylation, enrichment of specific histone marks (e.g. mono-methylation at histone H3 lysine 4 and acetylation of H3 lysine 27), as well as presence of a nucleosome-depleted region (Stadler et al. 2011; Thurman et al. 2012). While much work has focused on understanding the role of aberrant TF expression in leukemia, less is known about the role of the chromatin environment, and hence DNA methylation (Blattler and Farnham 2013), in modulating TF access to their cognate binding sites.

The occurrence of enhancer DNA hypermethylation and hematological malignancies upon *TET2* mutation challenges a widely held view that DNA methylation does not pose a challenge for TF binding (Thurman et al. 2012). Although several TFs have been demonstrated to bind methylated DNA and induce DNA hypomethylation (e.g. CTCF and REST) (Lienert et al. 2011; Stadler et al. 2011), many TFs show an inherent binding preference *in vitro* for motifs with either methylated or unmethylated CpG sites (Yin et al. 2017). As an illustration of this, global loss of DNA methylation in embryonic stem (ES) cells was shown to unmask previously inaccessible genomic binding sites for the TF NRF1 (Domcke et al. 2015). Moreover, DNA methylation and TET-mediated oxidation products potentially influence binding of epigenetic “readers” (e.g. Methyl-Binding Domain (MBD) proteins) (Song and Pfeifer 2016), histone variants and nucleosome remodeling enzymes (Conerly et al. 2010; Brunelle et al. 2015), and modify the physical properties of chromatin and DNA itself (Ngo et al. 2016). Thus, the dynamics of DNA methylation and TET-mediated cytosine oxidation at gene-regulatory elements may, under certain circumstances, modulate their function.

Despite advances in the identification of epigenetic changes in *TET2* mutated cells, the mechanisms by which aberrant DNA methylation promotes hematopoietic stem cell expansion and malignancy remain poorly understood. Here, we use genomic profiling to dissect the functional consequences of TET2 ablation in ES cells and *in vivo* models of CH/CCUS and AML. We find that TET2 binding and catalytic activity at enhancers facilitate recruitment of key hematopoietic TFs and we propose that this, mediates at least some of the phenotypic effects of *TET2* mutations in hematopoiesis.

## RESULTS

### TET2 binds regions of open chromatin with enhancer features in embryonic stem cells

Depletion of TET2 in ES and hematopoietic cells results in widespread changes in the DNA methylation landscape (Ko et al. 2010; Asmar et al. 2013; Hon et al. 2014; Rasmussen et al. 2015; Yamazaki et al. 2015). However, it still remains unclear which regions are directly targeted by TET2 as opposed to being secondary effects of its depletion. To address this, we developed and optimized reagents and protocols for genome-wide mapping of TET2 occupancy using chromatin immunoprecipitation (ChIP) in wildtype and epitope-tagged cell lines (see Supplemental methods and Fig. S1 for full detail).

We first determined the genomic binding regions of TET2 in ES cells using antibodies to TET2 (TET2-N) or to an epitope-tagged TET2 expressed from the endogenous *Tet2* locus (FLAG M2). TET2 binding sites were defined by enrichment over non-specific ChIP enriched regions identified in *Tet2* knockout cells or parental cells without FLAG-tagged TET2. In total, this analysis revealed 26,512 TET2-bound regions of which approximately one-third, referred to as “high-confidence” TET2 binding sites, were identified with both antibodies and had a stronger ChIP-seq signal (Fig 1A). Strikingly, the vast majority (93.4%) of these 8,262 sites were localized in regions of open chromatin defined by DNaseI hypersensitivity (DHS) (Fig. 1B and 1C).

**Figure 1:**
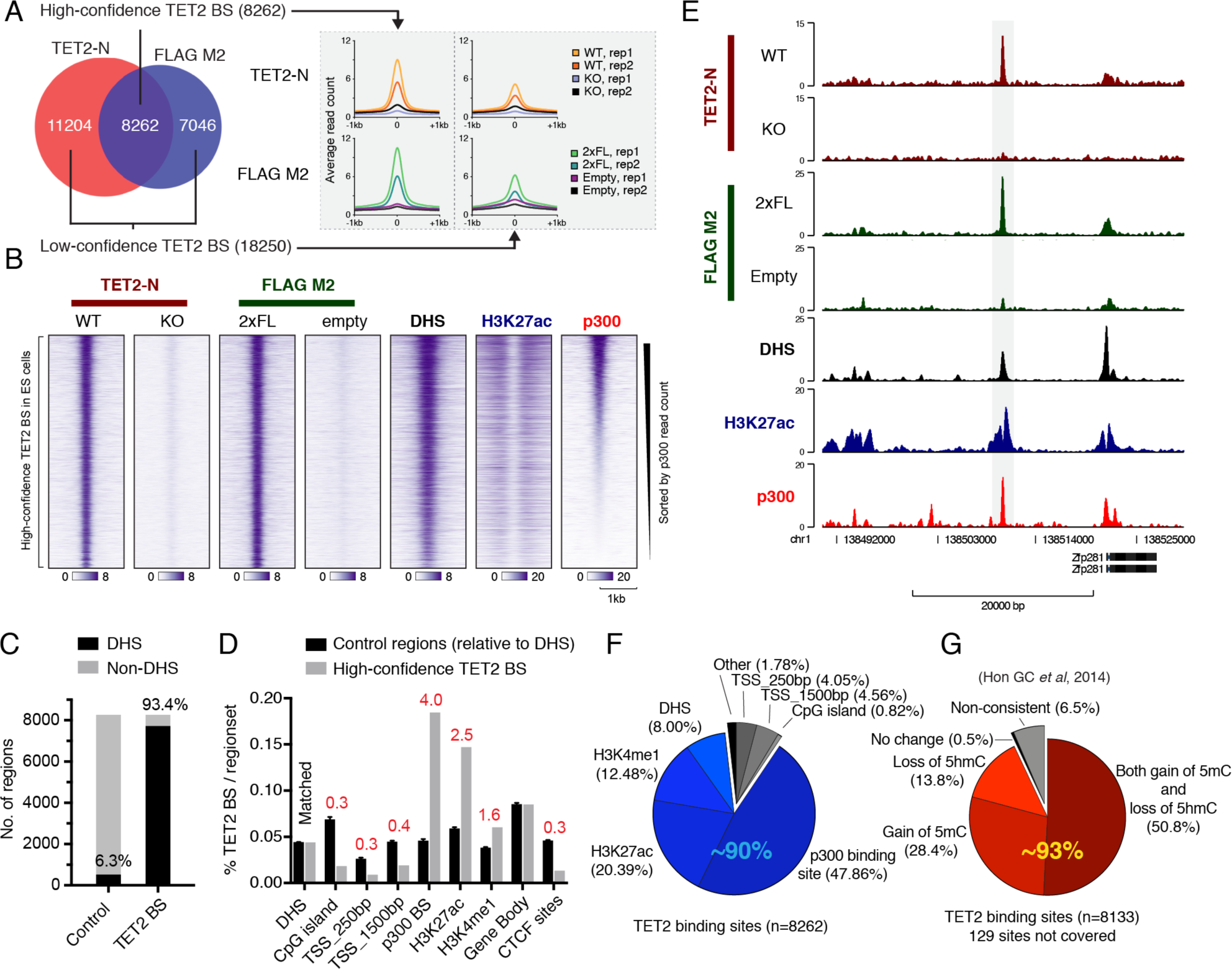
TET2 binds regions of open chromatin with enhancer features in embryonic stem cells. **(A)** Venn diagram showing overlap of called peaks in TET2-N or FLAG M2 ChlP-seq experiments (left panel) as well as average ChIP signal from replicate samples (Right panel). High- and low-confidence TET2 binding sites are defined, respectively, as regions showing evidence of TET2 binding in both peak sets (High) or only supported by a called positive region in one peak set (Low). **(B)** Heat maps of ChlP-seq signal from wildtype TET2 or TET2 with two copies of a FLAG tag (2xFL). Tracks of H3K27ac and p300 enrichment as well as regions of DHS in ES cells are also shown. The vertical axis contains all high-confidence TET2 binding sites defined in (A) sorted by decreasing p300 read counts. The horizontal axis is centered on TET2 peaks. **(C)** Histogram showing overlap of high-confidence TET2 binding sites with regions of DHS in ES cells. Random matched control regions (Control) were generated with same size, orientation and distance relative to gene bodies. **(D)** Histogram showing overlap as (C) but with random matched control regions relative to DHS. The percentage of CpG islands, proximal (+/-250bp) or distal (+/-1500bp) transcription start sites (TSS), p300 binding sites, H3K27ac and H3K4me1 enriched domains, Gene body and CTCF binding sites, overlapping TET2 binding sites are shown. Fold enrichment in each comparison are labeled in red. Overlap for control regions are plotted as mean +/-SD (n=5). **(E)** Representative ChlP-seq tracks showing a TET2-bound region upstream of the Zfp281 gene in ES cells. **(F)** Pie chart showing the distribution of high-confidence TET2 binding sites with respect to the indicated genomic regions. Each TET2 binding site are counted only once and excluded when moving clockwise from proximal promoters. Blue hues indicate fraction of TET2-bound regions with regulatory potential at promoter-distal sites (∼90% of all TET2-bound regions). **(G)** Pie chart as (F) showing the average change in DNA methylation (5mC and 5hmC) upon *Tet2* knockout in ES cells (Hon et al., 2014) in high-confidence TET2 binding sites and flanking regions (+/-250bp). Only CpG sites covered by >10 reads were included in the analysis. Each TET2 binding site are counted only once and excluded when moving clockwise. Red hues indicate fraction (∼93%) of TET2-bound regions showing DNA methylation changes consistent with loss of TET2 catalytic activity (gain of 5mC or loss of 5hmC, or both). Regions in **(B-F)** enriched for DHS, p300, H3K27ac, H3K4me1, and CTCF sites in ES cells were experimentally determined in the ENCODE project (ENCODE Project Consortium, 2011).

To understand whether TET2 is enriched in specific regions of open chromatin in the genome, we compared high-confidence TET2 binding sites to sets of random control regions (x5) matched for size, orientation, and distance to DHS sites. This showed a strong enrichment of TET2 high-confidence binding sites at DHS with enhancer features such as p300 binding (4-fold), H3K27ac (2.5-fold), and H3K4me1 (1.6-fold), whereas TET2 binding at promoters, CpG islands, and CTCF binding sites was depleted (∼0.3-fold each) (Fig 1D). This binding pattern is illustrated by a representative TET2-bound enhancer region 10kb upstream of the master ES regulator *Zfp281* (Fig. 1E). Notably, up to 90% of TET2 high confidence binding regions are associated with features of distal regulatory elements (p300, H3K27ac, H3K4me1, or DHS) and nearly half of TET2-bound regions overlap with promoter-distal p300 binding sites (Fig 1F). Chromatin occupancy of p300 is a hallmark of active enhancers and a biochemical interaction between p300 and TET2 has recently been reported (Heintzman et al. 2007; Zhang et al. 2017). Thus, our data are in agreement with a model in which TET2 can be directly recruited to a subset of its chromatin targets via a direct interaction with p300 (Zhang et al. 2017).

A previous study has examined the genome-wide changes in 5-methylcytosine (5mC) and 5-mhydroxymethylcytosine (5hmC) that occur upon *Tet2* knockout in ES cells (Hon et al. 2014). We used this dataset to identify epigenetic changes around (+/- 250bp) high-confidence TET2 binding sites. Genetic knockout of *Tet2* is likely to result in gain of 5mC (due to absence of DNA demethylation), loss of 5hmC (stable product of methylcytosine oxidation by TET2), or both, at sites of TET2 recruitment. Importantly, we found such changes (hyper-5mC, hypo-5hmC, or both) at the majority of high-confidence TET2 binding sites (∼93%) (Fig. 1G) thereby confirming that TET2 modifies the DNA methylation state. The analysis was repeated with the full set of 26,512 sites with evidence of TET2 binding (Fig. S2A). Within this larger and less stringent set of regions, we observed a modest enrichment of TET2 binding at CpG islands and gene promoters (Fig. S2B). However, in contrast to a pronounced DNA hypermethylation at sites co-bound by TET2 and p300, DNA methylation did not change at these CpG islands and promoters in *Tet2*-deficient ES cells (Fig S2C and S2D). This is consistent with the previously identified role of TET2 in maintaining a low level of DNA methylation at distal regulatory elements, whereas CpG islands and promoters are protected from DNA hypermethylation by additional mechanisms (Rasmussen and Helin 2016). It should be noted that most differentially methylated regions reported in *Tet2* knockout ES cells (over 60,000 hyper-DMRs and 130,000 hypo-DMRs) did not show detectable binding of TET2 in our dataset (Fig. S2E). This suggests either that TET2 functions in the absence of robust and persistent binding at these sites or that some of these methylation changes (especially hypo-DMRs) can occur as a consequence of secondary events to TET2 depletion. Taken together, our data show that TET2 binds to regions of open chromatin with enhancer features that undergo TET2-dependent DNA demethylation.

### Cell-type specific binding pattern of TET2 in myeloid hematopoietic cells versus ES cells

Next, we determined the genome-wide binding sites of TET2 in hematopoietic cells. To this end, we used myeloid cells immortalized by AML1-ETO with the potential for inducible deletion of *Tet2* (*Tet2*^*fl/fl*^; *AE; Rosa26*^*+*^/CreERT^2^ (Rasmussen et al. 2015)). Analysis of ChIP experiments using the TET2-N antibody identified 19706 regions significantly enriched over knockout control. Notably, only 7.4% of these regions were shared between ES cells and myeloid cells (Fig. 2A).

**Figure 2:**
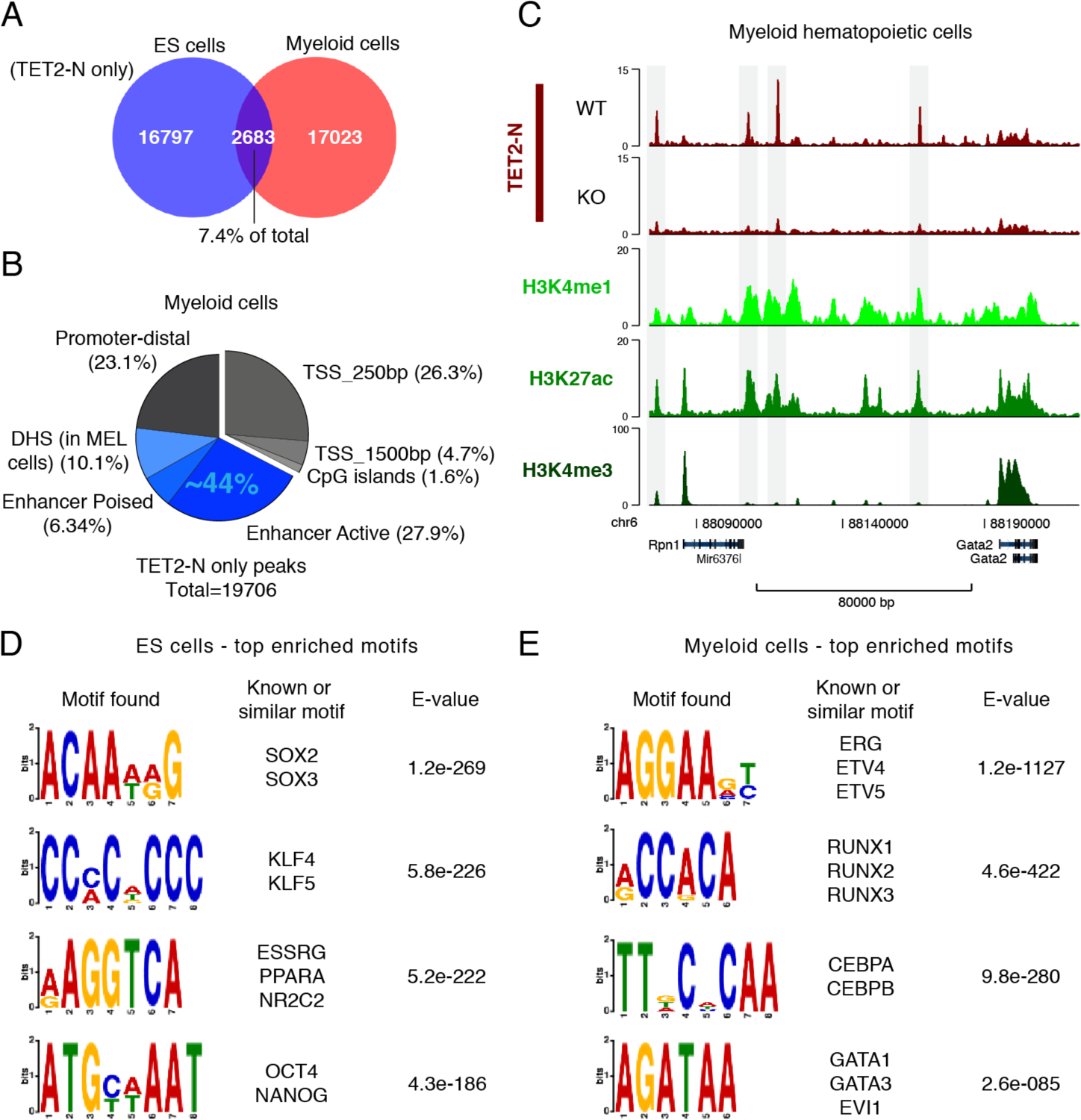
Cell-type specific binding pattern of TET2 in myeloid hematopoietic cells. **(A)** Venn diagram showing overlap of TET2 binding sites in ES cells and myeloid hematopoietic cells. Binding sites are defined by enrichment over non-specific ChIP enriched regions identified in Tet2 knockout cells using the TET2-N antibody. **(B)** Pie chart showing distribution of TET2-bound regions in myeloid hematopoietic cells. Each TET2 binding site are counted only once and excluded when moving clockwise from proximal promoters. Blue hues indicate fraction (∼44%) of TET2-bound regions with regulatory potential at promoter-distal sites based on H3K27 acetylation and H3K4 methylation in myeloid cells as well as DHS in the related mouse erythroleukemia (MEL) cell line. **(C)** Representative ChlP-seq tracks showing TET2-bound enhancer regions upstream of the Gata2 gene in myeloid hematopoietic cells. **(D)** List of top enriched sequence logos and their associated DNA-binding TFs identified in promoter-distal TET2-bound regions in ES cells. **(E)** as in (D), but for myeloid hematopoietic cells. Regions in (B) and (C) enriched for H3K27ac, H3K4me3, and H3K4me1 were experimentally determined in (Rasmussen et al., 2015) and DHS in MEL cells was downloaded from ENCODE project (ENCODE Project Consortium, 2011).

We then overlapped the TET2 myeloid binding sites with various genomic regions. Although binding of TET2 could be detected in a subset of promoters and CpG islands, the majority of binding sites was promoter-distal (67.4%) and nearly half (44.4%) associated with enhancer features (Fig. 2B). An illustrative example of this binding pattern is shown at enhancers upstream of the hematopoietic master regulator *Gata2* (Fig. 2C). Since promoter DNA methylation patterns are largely unaffected by depletion of TET2, we decided to focus our analysis on promoter-distal TET2 binding sites. Identification of enriched gene ontology (GO) terms by genomic regions enrichment of annotations tool (GREAT) revealed a striking separation by cell type. For example, the GO terms “stem cell maintenance” and “blastocyst formation” were enriched for TET2 binding sites in ES cells, whereas “immune system process” was highly significant for sites in myeloid cells (Fig. S2F and S2G). Next, we performed motif enrichment analysis of known DNA-binding TFs. The top enriched motifs belonged to master TFs of the respective cell types such as SOX2, KLF4, ESRRG, OCT4 and NANOG in ES cells and e.g. ERG, RUNX1, CEBPA, GATA1 in myeloid cells (Fig. 2D and 2E). Together, these results indicate that TET2 is recruited to chromatin in a highly cell-type specific manner and that TET2 binding at promoter-distal regulatory elements co-localizes with a wide range of predicted TF binding sites.

### Native hematopoiesis in aged *Tet2*-deficient animals as a model system of CH/CCUS

Mutations in *TET2* are frequently observed in individuals with CH/CCUS (Jaiswal et al. 2014; Genovese et al. 2014; Xie et al. 2014; Kwok et al. 2015; Hansen et al. 2016). These mutations arise in hematopoietic stem cells with multi-lineage differentiation potential and confer a competitive advantage that, over an extended period of time, allows the mutated clone to expand and renders terminal differentiation inefficient (Bowman et al. 2018). To recapitulate this in a mouse model, we induced hematopoietic-specific *Tet2* deletion in young adult mice and allowed them to age to 10 months (40 weeks). Aged *Tet2*-deficient mice had increased levels of Lin^-^Sca^+^cKit^+^ (LSK) cells in the bone marrow and spleen, mild splenomegaly, and a mild anemia in the peripheral blood (Fig S3A-S3D). Thus, this mouse model approximates hematopoietic features observed in aged individuals with CH/CCUS and allows for molecular profiling of hematopoietic stem cells in their native microenvironment.

It has been demonstrated that native hematopoiesis is predominantly sustained by multipotent progenitors (MPP) throughout the lifetime of the mouse (Sun et al. 2014; Busch et al. 2015). We therefore decided to perform genomic profiling on wildtype and *Tet2*-deficient MPPs as well as downstream myeloid-lineage progenitors (Granulocyte-Monocyte Progenitors, GMP) (Fig 3A and S3E). Analysis of differentially regulated genes in TET2 depleted cells revealed relatively few expression changes in MPPs (106 genes, *q*- value < 0.1), whereas GMPs had 8-fold higher number of up- and down-regulated genes (Fig S3F and Table S1). Yet, comparison of differentially expressed genes identified several important regulators of hematopoiesis to be aberrantly expressed in both cell types (e.g. Up: *Zbtb16, Runx1, Pdgfrb,* down: *Etv5, Gfi1*) (Fig. 3B). Interestingly, gene set enrichment analysis (GSEA) revealed a general downregulation in *Tet2*- deficient MPPs of gene signatures of translation, ribosome biogenesis and metabolism (Fig. 3C, S3G and Table S2). This is consistent with a report identifying lower metabolic rate and reduced ribosome biogenesis as hallmarks of preleukemic stem cells (Cai et al. 2015). Indeed, the total content of RNA, most of which is comprised of ribosomal RNA, was decreased in sorted *Tet2*-deficient MPPs (Fig. 3D).

**Figure 3:**
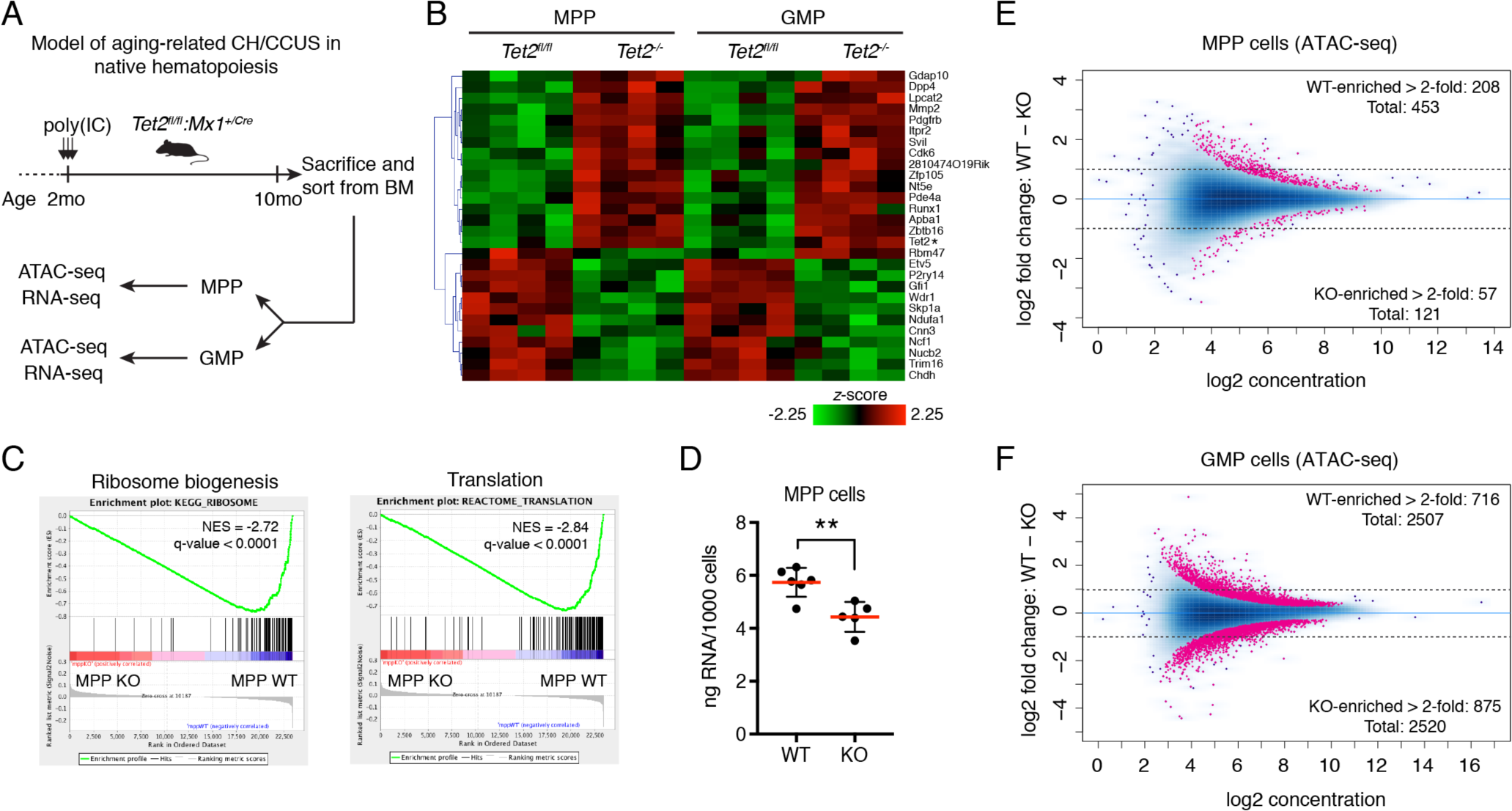
Native hematopoiesis in aged 7ef2-deficient animals as model system of CH/CCUS. **(A)** Diagram showing experimental setup to analyze native hematopoiesis in aged 7e/2-deficient animals. MPP, Multipotent progenitors. GMP, Granulocyte-Monocyte progenitors. **(B)** Heat map of variance-stabilized mRNA counts from RNA-seq in wildtype and *Te/2*-deficient cells (n=4). Only genes showing significant change (FDR < 0.1) in both MPPs and GMPs are shown (see Table S1 for full list of differentially expressed genes). Asterix denotes artefact of TET2 expression (only reads from N-terminal fragment of TET2 that is stably expressed after the catalytic domain in exon 11 has been deleted (Quivoron et al., 2011)). **(C)** Enrichment plots from GSEA showing decreased expression in Te/2-deficient MPPs of genes involved in ribosome biogenesis and translation (see top 20 wildtype enriched gene signatures in Fig. S3G and Table S2). **(D)** Scatter dot plot showing total RNA amount in MPPs sorted from aged animals with wildtype or 7ef2-deficient hematopoiesis in an independent experiment. Line and error bars represents mean +/-SD (n=5-6). **, P < 0.005 (unpaired two-tailed Student’s t-test). **(E)** Differentially accessible regions from ATAC-seq analysis of wildtype or 7ef2-deficient MPPs (n=4). Dots indicate regions with significant (FDR < 0.05) WT- or KO-enriched ATAC-seq signal. **(F)** as is (E), but for GMPs.

Profiling of chromatin accessibility by Assay for Transposase Accessible Chromatin and sequencing (ATAC-seq) (Buenrostro et al. 2013) revealed, analogous to the transcriptome analysis, relatively minor changes in MPPs (574 differential regions, q-value < 0.05). However, the majority of these regions (∼80%) were less accessible in *Tet2*-deficient cells - consistent with a direct effect of increased DNA methylation - and a considerable fraction (∼37%) overlapped with previously annotated enhancers in the blood lineages (Fig. 3E and S3H) (Lara-Astiaso et al. 2014). In contrast, we detected 10 times more differentially accessible regions in GMPs (5027 differential regions, q-value < 0.05) that were balanced between higher and lower chromatin accessibility (Fig. 3F and S4B) indicating that accumulated epigenetic dysregulation or associated secondary events (e.g. imbalance of differentiation-associated cytokines and growth factors) results in aberrant myeloid differentiation particularly as the cells progress towards GMPs. In summary, this shows that aged *Tet2*-deficient MPPs with features of preleukemic hematopoiesis are still closely related to the wildtype in terms of chromatin and transcriptional states and analysis of these can potentially unveil alterations directly related to loss of TET2 in hematopoietic stem cells.

### Differential TF analysis of chromatin accessibility reveals widespread changes in TF activity upon TET2 loss

Chromatin accessibility in gene regulatory regions can be used as a genome-wide measure of non-histone protein binding to DNA (Hesselberth et al. 2009). To assess whether a particular set of TFs is driving the changes we observed upon *Tet2* knockout, we developed a novel computational method, *diffTF* (Berest et al *in revision*), to assess “TF activity” on a genome-wide scale using profiles of chromatin accessibility by ATAC-seq. In this method we compare the accessibility changes at putative binding sites for each TF and compare this distribution to the background distribution of accessibility fold changes for all other TFs. If the putative binding sites of a TF are overall less open in the *Tet2* mutant we define this TF to be less “active” in the *Tet2* depleted condition and vice versa. Thus “TF activity” is here defined as the TF being associated with increased chromatin accessibility at its target sites.

Loss of TET2 leads to widespread and pleiotropic changes in TF activity in MPP as well as GMP cells (Fig. 4A and Fig. S4). In GMP cells, we could detect a signature of aberrant lineage differentiation characterized by increased activity of the IRF family of TFs (IRF8, IRF1 and IRF2) and decreased activity of TAL1 and the GATA family members (GATA1, −2 and −3). Conversely, MPP cells showed a significant increase in chromatin accessibility in predicted binding sites of the CCAAT/Enhancer binding protein family (C/EBP- alpha, -beta, -gamma, -delta, HLF) as well as CCCTC-binding factor (CTCF) protein. Interestingly, previous studies have suggested that C/EBP-beta has a slight preference for binding to motifs with a methylated cytosine (Mann et al. 2013; Zhu et al. 2016), thus potentially implicating TET2 as a factor that restricts C/EBP- beta binding to chromatin. In the other extreme, we observed a significant decrease in the activity of several TFs including homeodomain TFs CDX4, EVX1 and HOXC6, as well as the E proteins: E2-2 and ZEB1.

**Figure 4:**
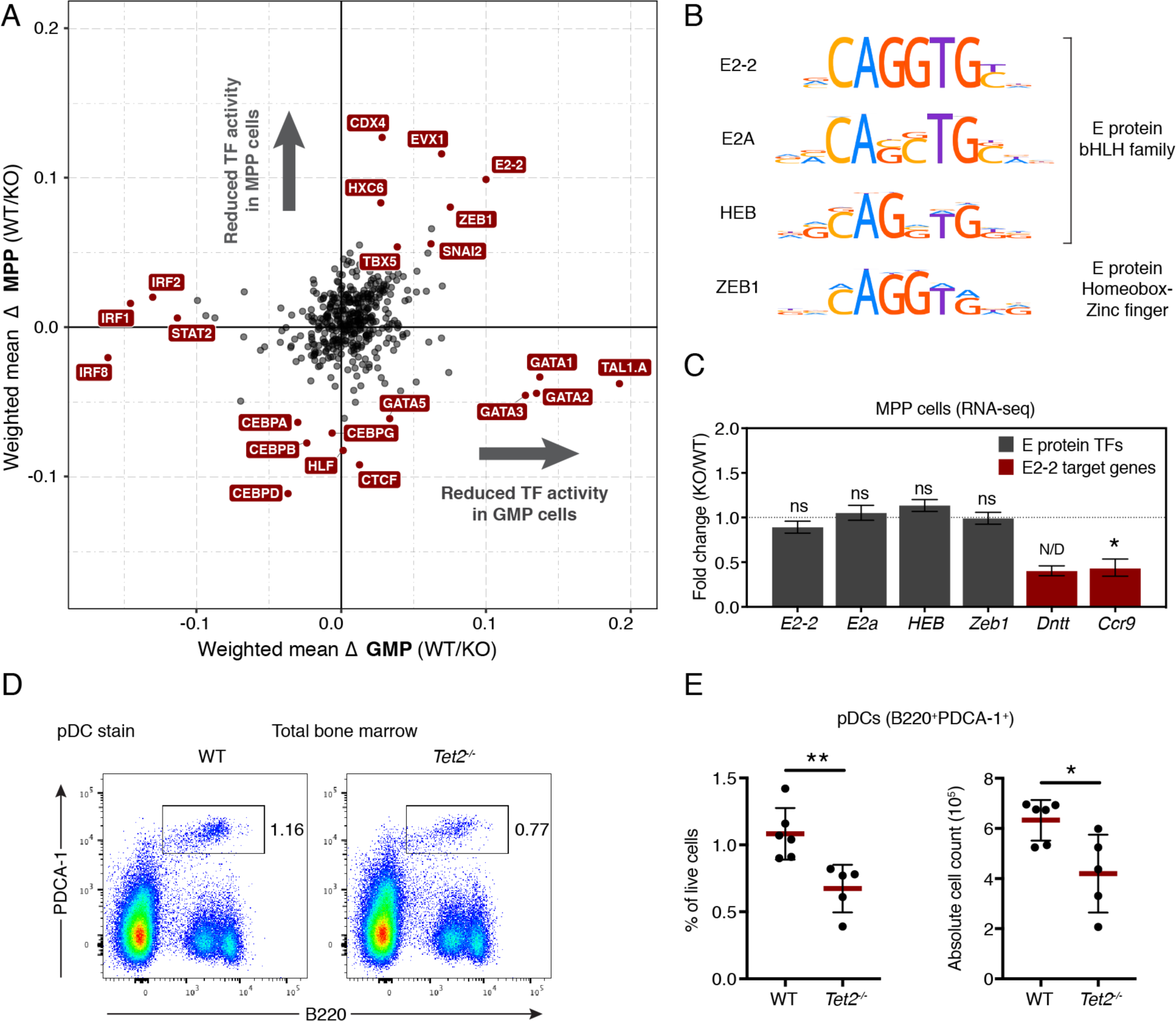
Differential TF analysis of chromatin accessibility reveals widespread changes in TF activity upon loss of TET2. **(A)** Scatter plot showing weighted mean diffTF values for 423 murine DNA binding motifs in MPPs (x-axis, n=4) or GMPs (y-axis, n=4). Positive and negative values indicate wildtype and KO-enriched activity, respectively. Red dots indicate TFs that pass a significance threshold (FDR < 0.01) and have a mean difference greater than +/- 0.05 (MPP) or +/- 0.1 (GMP) (See also Fig. S4 and S6A-D for full detail). **(B)** Sequence logos of the E protein bHLH family E2-2, E2A and HEB, as well as the related ZEB1 homeobox zinc-finger TF. **(C)** DESeq2 expression values from RNA-seq in MPPs of TFs in (B) as well as the two E2-2 target genes Dntt and Ccr9 (marked in red). Data is presented as mean +/- SEM (n=4). ns, not significant. N/D, not calculated due to outlier sample. \ P < 0.05 (q-value). **(D)** Representative FACS plots showing the population of plasmacytoid dendritic cells (pDC) in bone marrow of aged wildtype and Tef2-deficient animals. **(E)** Scatter dot plots showing the percentage (left) and absolute cell count (right) of pDCs in the bone marrow of aged wildtype and 7e/2-deficient animals. Line and error bars represent mean +/-SD (n=5-6). *, P < 0.05. **, P < 0.01 (unpaired two-tailed Student’s t-test).

To validate the biological significance of these changes in hematopoietic differentiation we focused on E2-2, whose activity showed a decrease upon *Tet2* knockout in both MPPs and GMPs (Fig. 4A and Fig. S4). E2-2 is a well-studied hematopoietic TF (reviewed in (Kee 2009)), and is a member of the basic Helix-Loop-Helix (bHLH) TF family whose *in vitro* DNA binding is inhibited by DNA methylation (Yin et al. 2017). In contrast to the loss of TF activity, we found that the mRNA levels of E2-2 as well as the closely related E proteins E2A, HEB and ZEB1 (Fig. 4B) were unchanged in *Tet2*-deficient MPPs (Fig. 4C). Mice with hematopoietic-specific knockout of one allele of *E2-2* display defects in differentiation of plasmacytoid dendritic cells (pDCs) and low expression of key pDC-associated genes (Cisse et al. 2008). We reasoned that a change in E2-2 activity should recapitulate these effects of reduced E2-2 expression. Consistent with this, we observed decreased expression of the target genes *Ccr9* and *Dntt* in *Tet2* knockout MPPs (Fig. 4C) and a partial impairment of pDC development in *Tet2*-deficient animals (Fig 4D and 4E).

Overall, these results suggest that the knockout of *Tet2* phenocopies the haploinsufficiency of E2-2 in hematopoietic stem cells by decreasing the binding activity of E2-2 rather than changing its expression. This would derail normal pDC differentiation in the bone marrow. Interestingly, frequent mutations of *TET2* have been found in patients with Blastic plasmacytoid dendritic cell neoplasm (BPDCN), a rare myeloid neoplasm characterized by proliferation of aberrant pDC precursor cells (Scourzic et al. 2015). Thus, TET2 may prevent BPDCN by keeping the binding sites of E2-2 accessible and thus maintaining the expression of E2-2-regulated genes.

### Attenuation of CDX4 activity in native hematopoiesis and AML with loss of TET2

Mutations of *TET2* are both implicated in the initiation and the maintenance of hematopoietic malignancies (Cimmino et al. 2017). Therefore, we set out to compare the *diffTF* signature of MPP cells and fully transformed leukemia with ablation of TET2. To do this, we generated a novel mouse model of human normal karyotype AML with mutations in *TET2* and the frequently co-occurring mutations *NPM1c* and *FLT3-ITD* (Fig. S5A). A combination of the oncogenic mutations *Npm1c* and *Flt3-ITD* in the mouse is sufficient to drive formation of AML (Mupo et al. 2013), however simultaneous deletion of *Tet2* significantly increased *in vitro* colony-forming capacity and accelerated the onset of disease (Fig. S5B and S5C). Furthermore, primary leukemic cells from moribund mice can be maintained and expanded *in vitro* and these cells maintain leukemogenicity even after extensive cell culture passaging (Fig S5D and S5E). Characterization of surface marker expression in the bulk *in vitro* culture revealed sustained proliferation of CD34^+^CD16/32^int^CD11b^-^Gr1^-^ cells, whereas more differentiated cells in the clonal hierarchy did not grow (Fig. S5F). Thus, we purified this leukemogenic precursor population (CD34^+^CD16/32^int^CD11b^-^Gr1^-^) from wildtype and *Tet2*-knockout cultures for analysis of chromatin accessibility by ATAC-seq.

*diffTF* analysis of AML cells with *Tet2* knockout versus wildtype identified decreased TF activity of members of the *HoxB* gene cluster including HOXB7 and HOXB8 as well as the TFs MITF and STAT6 (Fig. 5A). In addition, we detected a pronounced attenuation of the two homeodomain TFs CDX4 and EVX1. Importantly, the decrease of CDX4/EVX1 activity upon loss of TET2 was observed in both MPP cells and AML cells suggesting that TET2 supports TF binding at these sites in both normal and malignant hematopoiesis (Fig. 5A). Due to the similarity of their reported DNA binding motif, the effects of CDX4 and EVX1 are difficult to distinguish based on binding site predictions (Fig. 5B). To disentangle their effects, we performed quantitative RT-PCR and found similar expression levels of CDX4 mRNA in wildtype and *Tet2* deleted LSK cells, whereas expression of EVX1 was not detectable in either (Fig. 5C). In addition, we observed similar levels of CDX4 protein expression in wildtype and *Tet2-null* AML and ES cells (Fig. 5D and 5E). Finally, a role for CDX4 has been described in early hematopoietic development in the embryo as well as adult hematopoiesis (Bansal et al. 2006; Wang et al. 2008; Koo et al. 2010; Rawat et al. 2012; McKinney-Freeman et al. 2008), whereas EVX1 has yet to be implicated in these processes. Hence, we conclude that the reduced accessibility at predicted CDX4/EVX1 sites in hematopoietic cells most likely stems from loss of CDX4 binding upon loss of TET2. Reduction of CDX4 activity furthermore occurs predominantly at promoter-distal regions (Fig. S4A see “AML.ProDistal”) and is observed at sites of TET2 binding in AML cells (Fig. S4A see “AML.ChIP”). To assess the functional role of the altered TF activity, we analyzed the relationship between TF activity differences reported by *diffTF* and the differential expression pattern of predicted target genes in proximity to the TF binding sites using RNA-seq data from MPPs. Importantly, we observed an overall decrease of CDX4/EVX1 target gene expression thus confirming the role of CDX4 as an activator of gene expression (Fig. S6E).

**Figure 5:**
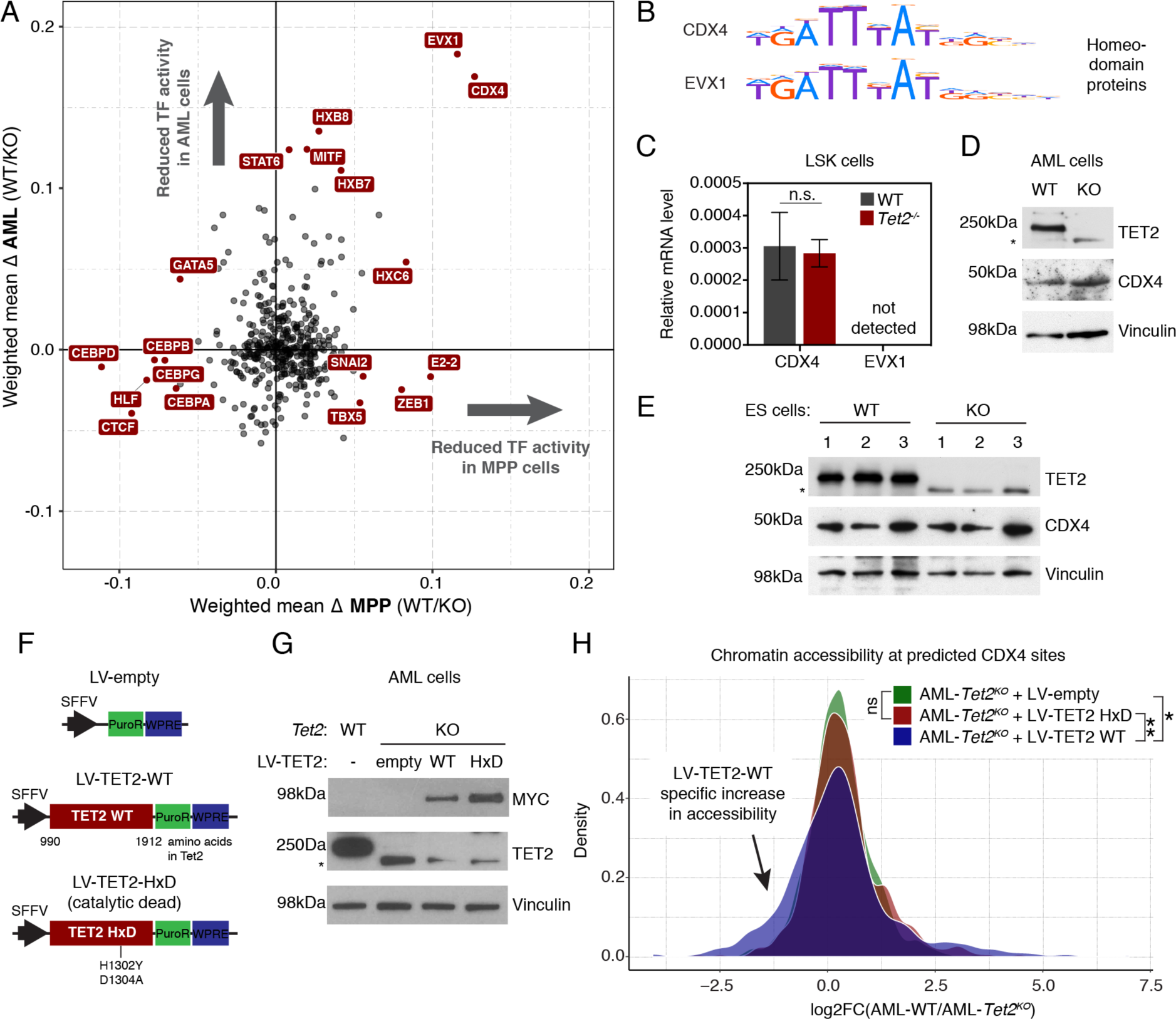
Attenuation of CDX4 activity in native hematopoiesis and AML with loss of TET2. **(A)** Scatter plot showing weighted mean diffTF values for 423 murine DNA binding motifs in AML cells (x-axis, n=3) or MPP cells (y-axis, n=4). Positive and negative values indicate wildtype and KO-enriched activity, respectively. Red dots indicate TFs that pass a significance threshold (FDR < 0.01) and a mean difference greater than +/- 0.05 (MPP) or +/- 0.1 (AML) (See also Fig. S4 and S6A-D for full detail). **(B)** Sequence logos for the DNA binding motif of the homeodomain TFs CDX4 and EVX1. **(C)** qRT-PCR analysis of CDX4 and EVX1 expression in LSK cells sorted from aged wildtype and 7e/2-deficient animals. The mean expression +/- SEM (n=5) relative to the housekeeping gene Hprt is shown, ns, non-significant. EVX1 mRNA was not detected. **(D)** Western blot for TET2 and CDX4 in AML cells with and without *Tet2* knockout. The loading control Vinculin is also shown. **(E)** Western blot as in (D), but in triplicate cultures of wildtype or 7e/2-deficient ES cells. **(F)** Schematic showing the lentiviral vectors (LV) used to overexpress myc-tagged TET2 catalytic domain or a catalytic inactive version (HxD, with amino acid substitutions H1302Y and D1304A). **(G)** Western blot (right panel) for MYC, TET2 and Vinculin loading control in AML cells with *Tet2* knockout and transduced with the indicated vectors**. (H)** Distribution plot showing log2 fold changes of chromatin accessibility at predicted CDX4 binding sites in AML-*Tet2*KO cells transduced with empty vector, TET2-WT or TET2-HxD (catalytic dead). The emergence of a left tail in the distribution indicates subset of sites with increased accessibility upon expression of TET2-WT, but not TET2-HxD. ns, not significant. *, P < 0.05. **, P < 0.005 (Unpaired two-tailed Mann-Whitney test). Asterix in **(D-F)** denotes the N-terminal fragment of TET2 that is stably expressed after the catalytic domain in exon 11 has been deleted (Quivoron et al., 2011).

To investigate whether chromatin recruitment of CDX4 is promoted by catalytic activity-dependent or -independent functions of TET2, we transduced AML cells with lentiviruses (LVs) expressing wildtype and a catalytic dead (HxD) version of the TET2 catalytic domain (Fig. 5F). We then quantified chromatin accessibility in predicted CDX4 binding sites by ATAC-seq. Despite higher expression of TET2-HxD compared to TET2-WT in the *Tet2* knockout cells (Fig. 5G), only TET2-WT was able to partially rescue CDX4 binding as indicated by increased chromatin accessibility at a subset of predicted binding sites (left tail) (Fig. 5H). This was furthermore confirmed by an overall increase in chromatin accessibility in cells expressing TET2-WT compared to cells expressing TET2-HxD (Mann-Whitney non-parametric, *P* = 0.0024) (Fig. 5H). Thus, we conclude that the catalytic activity of TET2 is necessary to sustain chromatin accessibility at a subset of predicted CDX4 binding sites, and that this, most likely, occurs by modulation of DNA methylation in the local chromatin environment to support CDX4 chromatin binding.

### Decreased CDX4 chromatin occupancy in *Tet2* knockout ES cells

The lack of ChIP-grade antibodies for CDX4 makes it difficult to directly assess its chromatin occupancy in hematopoietic cells. This prompted us to investigate *Tet2*-dependent CDX4 chromatin binding in ES cells, a cell type with a well-described role for CDX4 in promoting hematopoietic differentiation (Wang et al. 2008; McKinney-Freeman et al. 2008). We engineered a mouse ES cell line on a *Tet2*^*fl/fl*^; *Rosa26-Cre*^*+*^/ERT^2^ genetic background with stable expression of murine CDX4 with two copies of the FLAG tag (2xFL). This enables analysis of CDX4 chromatin occupancy before and after inducible deletion of *Tet2* by addition of 4-hydroxytamoxifen (4-OHT) (Fig. 6A). ChIP-seq analysis identified a total of 13,977 enriched regions in 2xFL-CDX4 expressing cells compared to the parental cells without 2xFL-CDX4 expression (Fig. 6B). As a quality control, we analysed DNA sequences at CDX4 enriched regions by TF DNA-binding motif enrichment analysis. The top enriched motif (E-value = 1.1e-529) was highly similar to the DNA binding motif of the ParaHox class of TFs (Cdx1, Cdx2, and Cdx4), thus providing an independent validation of the ChIP results (Fig. S6F).

**Figure 6:**
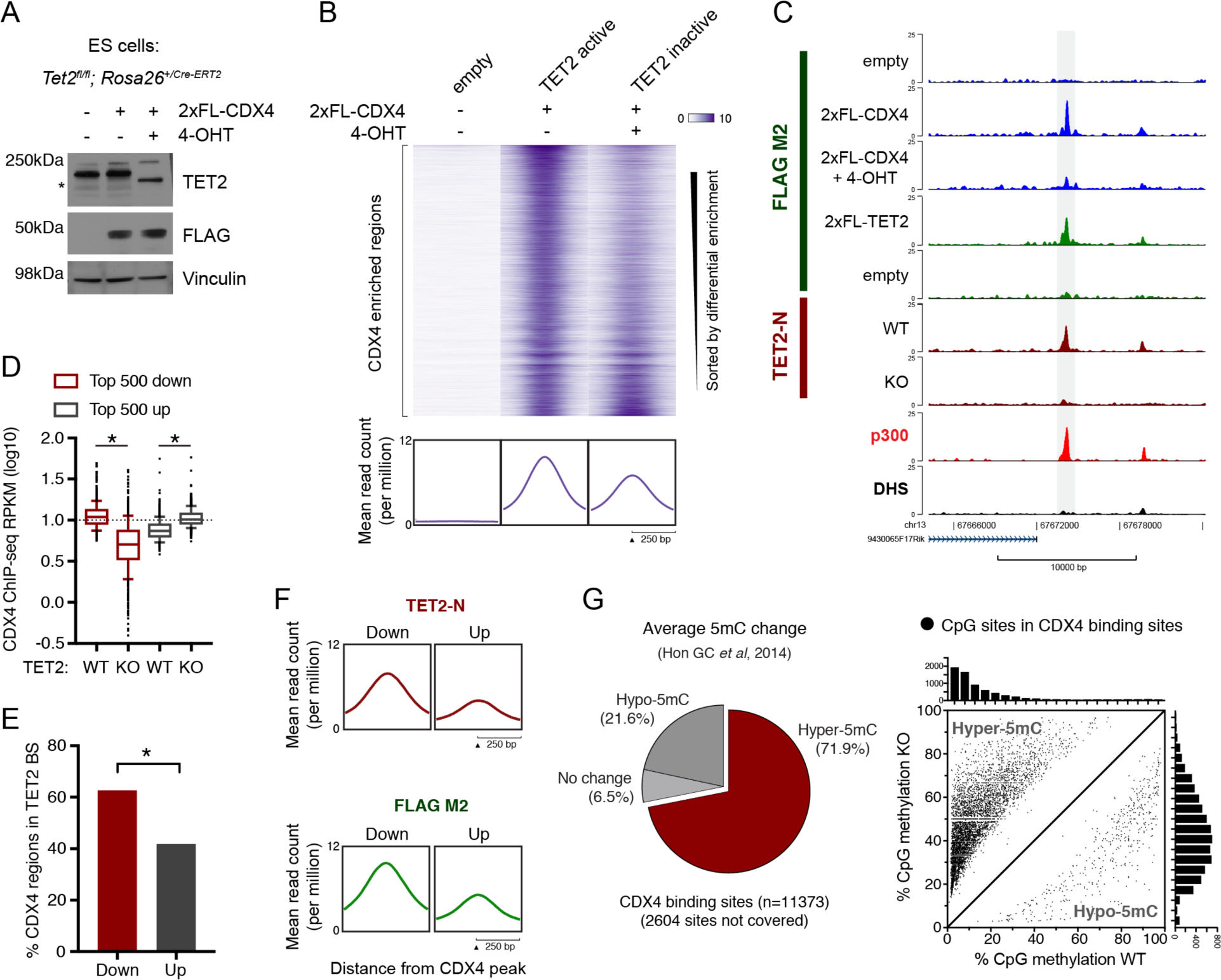
Decreased CDX4 chromatin occupancy in Tet2-deficient ES cells. **(A)** Western blot for 2xFL-tagged CDX4, TET2, and Vinculin loading control in *Tet2^a/n^*; *Rosa26*’^CreERT^*- embryonic stem cells. Treatment with 4-OHT induces Cre-mediated deletion of full-length TET2. Asterix denotes the N-terminal fragment of TET2 that is stably expressed after the catalytic domain in exon 11 has been deleted (Quivoron et al., 2011). **(B)** Heat maps of CDX4 ChlP-seq signal in ES cells with and without stable expression of 2xFL-CDX4 and inducible deletion of *Tet2*. The vertical axis contains all CDX4-enriched regions and is ordered by decreasing differential enrichment in Te/2-deficient cells versus wildtype cells expressing 2xFL-CDX4. The mean CDX4 ChlP-seq signal for each cell line is shown below. **(C)** Representative ChlP-seq tracks showing loss of CDX4 ChlP-seq signal upon 4-OHT induced deletion of *Tet2* ai a region co-bound by CDX4, *TET2* and p300 in ES cells. **(D)** Box plot showing log 10 of Reads Per Kilobase Million (RPKM) in top 500 CDX4 sites with decreased reads (down) or increased reads (up) upon ablation of *Tet2*. Box whiskers represents 10-90th percentile. \ P < 0.0001 (paired two-tailed Student’s t-test). **(E)** Histogram indicating the subset of CDX4 binding sites as in (D) that overlap with a called TET2 peak (all 26,512 peaks) in ES cells. \ P < 0.0001 (Fisher’s exact test). **(F)** Plots showing the average TET2 ChlP-seq signal in the subset of CDX4 binding sites as in (D and E). TET2 is robustly associated with CDX4 sites that show a decrease in chromatin occupancy upon ablation of *Tet2* (down) compared to control regions (up) using both TET2-N (upper panel) and FLAG M2 epitopes (lower panel). **(G)** Pie chart (left panel) showing the average change in DNA methylation (5mC) upon *Tet2* knockout in ES cells in CDX4 binding sites +/- 250bp. Only CpG sites covered by >10 reads were included in the analysis. Scatter plot (right panel) showing individual CpGs in CDX4 binding sites with significantly (q-value < 0.05, min. diff. 10%) altered DNA methylation upon knockout of *Tet2*\r\ ES cells.

Next, we evaluated the ChIP signal within CDX4 peaks after *Tet2* deletion. Although the effect on CDX4 binding sites was highly locus-specific, we found an overall loss of CDX4 chromatin occupancy upon *Tet2* knockout, and quantification revealed a decrease in normalized read counts at a majority of CDX4-bound regions (∼83%) (Fig. 6B). An illustrative example of loss of CDX4 binding is shown for an enhancer region cobound by CDX4, TET2 and p300 in ES cells (Fig. 6C). To test for a direct effect of TET2, we selected the top and bottom 500 regions with the strongest loss and gain of CDX4 binding, respectively (Fig. 6D), and overlapped these with TET2 binding sites in ES cells. Importantly, we found a significantly higher overlap of TET2-positive regions (Fig. 6E) as well as normalized read counts (Fig. 6F) in regions with loss of CDX4 binding compared to the control (Up) region set. Finally, assessment of DNA methylation changes using a previously published genome-wide analysis of DNA methylation in *Tet2* knockout ES cells (Hon et al. 2014), demonstrated that the majority of CDX4 binding sites are DNA hypermethylated upon ablation of TET2 (Fig. 6G). Together with the *diffTF* analysis, these results provide strong evidence that CDX4 chromatin occupancy is adversely affected by *Tet2* knockout likely through an increase in DNA methylation at CDX4 binding sites.

## DISCUSSION

While DNA methylation landscapes in *TET2*-mutated cells have been extensively characterized (Ko et al. 2010; Asmar et al. 2013; Hon et al. 2014; Rasmussen et al. 2015; Yamazaki et al. 2015), much less is known about the functional consequences that ultimately drive hematopoietic stem cell expansion and malignancy. Here, we resolve the inconsistency between regions reported to be bound by TET2 (Chen et al. 2013; Deplus et al. 2013; Peng et al. 2016) and those affected by TET2 catalytic function. We show that TET2 is predominantly recruited to promoter-distal regions of open chromatin, including enhancers. Therefore, it is likely that it is epigenetic perturbation of these elements that results in dysregulated hematopoiesis of *TET2*-mutated stem cells, possibly through pleiotropic effects on many genes that together derail homeostasis and differentiation. Interestingly, a recent study has reported a direct biochemical interaction between the histone acetyltransferase p300 and TET2 (Zhang et al. 2017). In ES cells, we find that approximately half of high-confidence TET2 binding sites co-localize with p300-enriched regions (Fig. 1A) and nearly half of all p300-enriched regions show evidence of TET2 binding (Fig. S2B). Taken together these results may suggest that p300 is involved in recruiting TET2 to chromatin through direct protein-protein interactions. However, a direct validation of this, as well as a comprehensive analysis of the role of additional potential recruiters, such as WT1 (Rampal et al. 2014; Wang et al. 2015), remain an area of active investigation.

To dissect the impact of TET2 loss in hematopoiesis, we investigated genome-wide changes in enhancer function using chromatin accessibility as a measure of activity in a native chromatin context. We report widespread changes, of which a majority of differentially open regions in all tested cell types, except GMP, were less accessible upon ablation of TET2 (Fig. S4B). This is consistent with a recent study showing that most DNA methylation-sensitive regulatory elements (∼88%) are inhibited by DNA methylation (Lea et al. 2017). To perform an unbiased analysis of TF signatures that were globally enriched or depleted, we developed a novel computational tool, *diffTF*, to assess differential TF binding from ATAC-seq profiles (Berest et al, *in revision*). Importantly, we could detect signatures of aberrant activity of several hematopoietic TFs in model systems of CH/CCUS and AML (Fig. S4A). It should be noted that additional TF binding events that were not identified in this study may also require TET2 activity; however, such TFs may not show strong global differences or are not detected from analysis of chromatin accessibility alone (e.g. because they lack a known binding motif).

We found decreased activity of the regulator of hematopoietic development, CDX4 (Bansal et al. 2006; Wang et al. 2008; Koo et al. 2010; Rawat et al. 2012; McKinney-Freeman et al. 2008), in the absence of changes in CDX4 mRNA or protein levels, in both native and malignant TET2-deficient hematopoiesis. Thus, this suggests that TET2 promotes CDX4 chromatin binding, and hence CDX4-driven gene expression. In support of this, we could detect a decreased expression of CDX4/EVX1 target genes in MPP cells upon ablation of TET2 (Fig. S6E). In addition, we show that the catalytic activity of TET2 is required for CDX4 TF activity and demonstrate that loss of CDX4 chromatin binding in *TET2*-null ES cells coincides with increased DNA methylation. It should be noted that the functional outcome of a DNA hypermethylation event caused by impaired TET2 function is likely to be strongly influenced by locus-specific factors such as CpG density and positioning, chromosomal conformation, properties of bound chromatin factors as well as redundant and/or combinatorial enhancer function. Accordingly, only a small fraction of predicted CDX4 binding sites (∼2.5%) contained a CG dinucleotide in their motif. This suggests that it is aggregate remodeling of the chromatin environment in and around the binding site, and not only direct DNA methylation at the motif, that disfavors CDX4 binding. Thus, our results support a model by which TET2 recruitment and catalytic functions are necessary to shape the chromatin structure for permissive TF binding at a majority of CDX4-bound enhancers in hematopoietic cells. The precise molecular events that culminate in loss of CDX4 binding at these sites remain to be determined.

Frequent mutations in the DNA methylation machinery have been found in patients suffering from hematological diseases as well as solid cancers (Rasmussen and Helin 2016). Yet, whether DNA methylation is causally involved in shaping gene expression patterns, rather than passively mirroring transcriptional states, is still a matter of debate (Schübeler 2015). Here, we present data in support of a role of TET2-mediated cytosine modifications at enhancers to facilitate TF binding and fidelity of target gene expression in hematopoietic cells. Rather than being an obligate activator, TET2 functions to reinforce binding of some TFs and contributes in this manner to enhancer activity and gene expression. Analysis of chromatin maturation after replication fork passage has indicated that open chromatin at cell-type specific enhancers are only slowly re-established through competition of TFs with newly deposited nucleosomes (Ramachandran and Henikoff 2016). Thus, cells that naturally divide rapidly and must undergo coordinated cell state transitions (e.g. hematopoietic stem and progenitor cells) may be particularly sensitive to epigenetic disturbances of TF binding kinetics. Comprehensive analysis of enhancer function in *TET2*-mutated hematopoiesis will foster a greater understanding of the role of epigenetic dysregulation in disease and may lead to discoveries of potential clinical significance.

## METHODS

Full description of methods and reagents used in this study can be found in the supplemental methods section.

## DATA ACCESS

The accession number for all ChIP-seq, ATAC-seq and RNA-seq reported in this study is GEO: GSE112974

## ACKNOWLEDGMENTS

We thank members of the Helin laboratory for advice and discussion. K.D.R. was supported by a postdoctoral fellowship from the Danish Medical Research Council (FSS 1333-00120B), K.N. was supported by Program for Advancing Strategic International Networks to Accelerate the Circulation of Talented Researchers, JSPS (S2704), L.S-C. was supported by a fellowship from EU (H2020-MSCA-IF-2017-796341). The work in the Helin laboratory was supported by grants to K.H. from The European Research Council (294666_DNAMET), the Danish Cancer Society, the Danish National Research Foundation (DNRF82), and through a center grant from the Novo Nordisk Foundation (NNF17CC0027852)).

## AUTHOR CONTRIBUTIONS

K.D.R designed the study, performed majority of experiments, analysed data, and wrote the first draft of the manuscript. I.B. and J.Z. provided genomic data analysis and interpretation (ATAC-seq, RNA-seq, ChIP-seq) and designed the *diffTF* software. S.K, K.N., L.S-C., and M.T.P. performed experiments. G.S.V. and J.C. provided essential reagents and intellectual input. K.H and J.Z. designed and supervised the study, analyzed data, secured funding and wrote the manuscript with K.D.R.

## DISCLOSURE DECLARATION

The authors declare non-conflicting interests.

## REFERENCES

An J, González-Avalos E, Chawla A, Jeong M, López-Moyado IF, Li W, Goodell MA, Chavez L, Ko M, Rao A. 2015. Acute loss of TET function results in aggressive myeloid cancer in mice. Nat Commun 6: 10071.

Asmar F, Punj V, Christensen J, Pedersen MT, Pedersen A, Nielsen AB, Hother C, Ralfkiaer U, Brown P, Ralfkiaer E, et al. 2013. Genome-wide profiling identifies a DNA methylation signature that associates with TET2 mutations in diffuse large B-cell lymphoma. Haematologica 98: 1912–1920.

Bansal D, Scholl C, Fröhling S, McDowell E, Lee BH, Döhner K, Ernst P, Davidson AJ, Daley GQ, Zon LI, et al. 2006. Cdx4 dysregulates Hox gene expression and generates acute myeloid leukemia alone and in cooperation with Meis1a in a murine model. Proc Natl Acad Sci USA 103: 16924–16929.

Blattler A, Farnham PJ. 2013. Cross-talk between site-specific transcription factors and DNA methylation states. J Biol Chem 288: 34287–34294.

Bowman RL, Busque L, Levine RL. 2018. Clonal Hematopoiesis and Evolution to Hematopoietic Malignancies. Cell Stem Cell 22: 157–170.

Brunelle M, Nordell Markovits A, Rodrigue S, Lupien M, Jacques P-É, Gévry N. 2015. The histone variant H2A.Z is an important regulator of enhancer activity. Nucleic Acids Res 43: 9742–9756.

Buenrostro JD, Giresi PG, Zaba LC, Chang HY, Greenleaf WJ. 2013. Transposition of native chromatin for fast and sensitive epigenomic profiling of open chromatin, DNA-binding proteins and nucleosome position. Nat Methods 10: 1213–1218.

Busch K, Klapproth K, Barile M, Flossdorf M, Holland-Letz T, Schlenner SM, Reth M, Höfer T, Rodewald H-R. 2015. Fundamental properties of unperturbed haematopoiesis from stem cells in vivo. Nature 518: 542–546.

Cai X, Gao L, Teng L, Ge J, Oo ZM, Kumar AR, Gilliland DG, Mason PJ, Tan K, Speck NA. 2015. Runx1 Deficiency Decreases Ribosome Biogenesis and Confers Stress Resistance to Hematopoietic Stem and Progenitor Cells. Cell Stem Cell 17: 165–177.

Chen Q, Chen Y, Bian C, Fujiki R, Yu X. 2013. TET2 promotes histone O-GlcNAcylation during gene transcription. Nature 493: 561–564.

Cimmino L, Dolgalev I, Wang Y, Yoshimi A, Martin GH, Wang J, Ng V, Xia B, Witkowski MT, Mitchell-Flack M, et al. 2017. Restoration of TET2 Function Blocks Aberrant Self-Renewal and Leukemia Progression. Cell 170: 1079–1095.e20.

Cisse B, Caton ML, Lehner M, Maeda T, Scheu S, Locksley R, Holmberg D, Zweier C, Hollander den NS, Kant SG, et al. 2008. Transcription factor E2-2 is an essential and specific regulator of plasmacytoid dendritic cell development. Cell 135: 37–48.

Conerly ML, Teves SS, Diolaiti D, Ulrich M, Eisenman RN, Henikoff S. 2010. Changes in H2A.Z occupancy and DNA methylation during B-cell lymphomagenesis. Genome Res 20: 1383–1390.

Deplus R, Delatte B, Schwinn MK, Defrance M, Méndez J, Murphy N, Dawson MA, Volkmar M, Putmans P, Calonne E, et al. 2013. TET2 and TET3 regulate GlcNAcylation and H3K4 methylation through OGT and SET1/COMPASS. EMBO J 32: 645–655.

Domcke S, Bardet AF, Adrian Ginno P, Hartl D, Burger L, Schübeler D. 2015. Competition between DNA methylation and transcription factors determines binding of NRF1. Nature 528: 575–579.

ENCODE Project Consortium. 2011. A user’s guide to the encyclopedia of DNA elements (ENCODE). ed. P.B. Becker. PLoS Biol 9: e1001046.

Genovese G, Kähler AK, Handsaker RE, Lindberg J, Rose SA, Bakhoum SF, Chambert K, Mick E, Neale BM, Fromer M, et al. 2014. Clonal hematopoiesis and blood-cancer risk inferred from blood DNA sequence. N Engl J Med 371: 2477–2487.

Hansen JW, Westman MK, Sjö LD, Saft L, Kristensen LS, Ørskov AD, Treppendahl M, Andersen MK, Grønbæk K. 2016. Mutations in idiopathic cytopenia of undetermined significance assist diagnostics and correlate to dysplastic changes. Am J Hematol 91: 1234–1238.

Heintzman ND, Stuart RK, Hon G, Fu Y, Ching CW, Hawkins RD, Barrera LO, Van Calcar S, Qu C, Ching KA, et al. 2007. Distinct and predictive chromatin signatures of transcriptional promoters and enhancers in the human genome. Nat Genet 39: 311–318.

Hesselberth JR, Chen X, Zhang Z, Sabo PJ, Sandstrom R, Reynolds AP, Thurman RE, Neph S, Kuehn MS, Noble WS, et al. 2009. Global mapping of protein-DNA interactions in vivo by digital genomic footprinting. Nat Methods 6: 283–289.

Hon GC, Song C-X, Du T, Jin F, Selvaraj S, Lee AY, Yen C-A, Ye Z, Mao S-Q, Wang B-A, et al. 2014. 5mC oxidation by Tet2 modulates enhancer activity and timing of transcriptome reprogramming during differentiation. Mol Cell 56: 286–297.

Jaiswal S, Fontanillas P, Flannick J, Manning A, Grauman PV, Mar BG, Lindsley RC, Mermel CH, Burtt N, Chavez A, et al. 2014. Age-related clonal hematopoiesis associated with adverse outcomes. N Engl J Med 371: 2488–2498.

Kee BL. 2009. E and ID proteins branch out. Nat Rev Immunol 9: 175–184.

Ko M, Huang Y, Jankowska AM, Pape UJ, Tahiliani M, Bandukwala HS, An J, Lamperti ED, Koh KP, Ganetzky R, et al. 2010. Impaired hydroxylation of 5-methylcytosine in myeloid cancers with mutant TET2. Nature 468: 839–843.

Koo S, Huntly BJ, Wang Y, Chen J, Brumme K, Ball B, McKinney-Freeman SL, Yabuuchi A, Scholl C, Bansal D, et al. 2010. Cdx4 is dispensable for murine adult hematopoietic stem cells but promotes MLL-AF9-mediated leukemogenesis. Haematologica 95: 1642–1650.

Kwok B, Hall JM, Witte JS, Xu Y, Reddy P, Lin K, Flamholz R, Dabbas B, Yung A, Al-Hafidh J, et al. 2015. MDS-associated somatic mutations and clonal hematopoiesis are common in idiopathic cytopenias of undetermined significance. Blood 126: 2355–2361.

Lara-Astiaso D, Weiner A, Lorenzo-Vivas E, Zaretsky I, Jaitin DA, David E, Keren-Shaul H, Mildner A, Winter D, Jung S, et al. 2014. Immunogenetics. Chromatin state dynamics during blood formation. Science 345: 943–949.

Lea AJ, Vockley CM, Johnston RA, Del Carpio CA, Barreiro LB, Reddy TE, Tung J. 2017. Genome-wide quantification of the effects of DNA methylation on human gene regulation. bioRxiv 146829.

Lienert F, Wirbelauer C, Som I, Dean A, Mohn F, Schübeler D. 2011. Identification of genetic elements that autonomously determine DNA methylation states. Nat Genet 43: 1091–1097.

Lu F, Liu Y, Jiang L, Yamaguchi S, Zhang Y. 2014. Role of Tet proteins in enhancer activity and telomere elongation. Genes Dev 28: 2103–2119.

Mahé EA, Madigou T, Sérandour AA, Bizot M, Avner S, Chalmel F, Palierne G, Métivier R, Salbert G. 2017. Cytosine modifications modulate the chromatin architecture of transcriptional enhancers. Genome Res 27: 947–958.

Mann IK, Chatterjee R, Zhao J, He X, Weirauch MT, Hughes TR, Vinson C. 2013. CG methylated microarrays identify a novel methylated sequence bound by the CEBPB|ATF4 heterodimer that is active in vivo. Genome Res 23: 988–997.

McKinney-Freeman SL, Lengerke C, Jang I-H, Schmitt S, Wang Y, Philitas M, Shea J, Daley GQ. 2008. Modulation of murine embryonic stem cell-derived CD41+c-kit+ hematopoietic progenitors by ectopic expression of Cdx genes. Blood 111: 4944–4953.

Mupo A, Celani L, Dovey O, Cooper JL, Grove C, Rad R, Sportoletti P, Falini B, Bradley A, Vassiliou GS. 2013. A powerful molecular synergy between mutant Nucleophosmin and Flt3-ITD drives acute myeloid leukemia in mice. Leukemia 27: 1917–1920.

Ngo TTM, Yoo J, Dai Q, Zhang Q, He C, Aksimentiev A, Ha T. 2016. Effects of cytosine modifications on DNA flexibility and nucleosome mechanical stability. Nat Commun 7: 10813.

Peng L, Li Y, Xi Y, Li W, Li J, Lv R, Zhang L, Zou Q, Dong S, Luo H, et al. 2016. MBD3L2 promotes Tet2 enzymatic activity for mediating 5-methylcytosine oxidation. J Cell Sci 129: 1059–1071.

Quivoron C, Couronné L, Valle Della V, Lopez CK, Plo I, Wagner-Ballon O, Do Cruzeiro M, Delhommeau F, Arnulf B, Stern M-H, et al. 2011. TET2 inactivation results in pleiotropic hematopoietic abnormalities in mouse and is a recurrent event during human lymphomagenesis. Cancer Cell 20: 25–38.

Ramachandran S, Henikoff S. 2016. Transcriptional Regulators Compete with Nucleosomes Post-replication. Cell 165: 580–592.

Rampal R, Alkalin A, Madzo J, Vasanthakumar A, Pronier E, Patel J, Li Y, Ahn J, Abdel-Wahab O, Shih A, et al. 2014. DNA hydroxymethylation profiling reveals that WT1 mutations result in loss of TET2 function in acute myeloid leukemia. Cell Rep 9: 1841–1855.

Rasmussen KD, Helin K. 2016. Role of TET enzymes in DNA methylation, development, and cancer. Genes Dev 30: 733–750.

Rasmussen KD, Jia G, Johansen JV, Pedersen MT, Rapin N, Bagger FO, Porse BT, Bernard OA, Christensen J, Helin K. 2015. Loss of TET2 in hematopoietic cells leads to DNA hypermethylation of active enhancers and induction of leukemogenesis. Genes Dev 29: 910–922.

Rawat VPS, Humphries RK, Buske C. 2012. Beyond Hox: the role of ParaHox genes in normal and malignant hematopoiesis. Blood 120: 519–527.

Schübeler D. 2015. Function and information content of DNA methylation. Nature 517: 321–326.

Scourzic L, Mouly E, Bernard OA. 2015. TET proteins and the control of cytosine demethylation in cancer. Genome Med 7: 9.

Song J, Pfeifer GP. 2016. Are there specific readers of oxidized 5-methylcytosine bases? Bioessays 38: 1038–1047.

Stadler MB, Murr R, Burger L, Ivanek R, Lienert F, Schöler A, van Nimwegen E, Wirbelauer C, Oakeley EJ, Gaidatzis D, et al. 2011. DNA-binding factors shape the mouse methylome at distal regulatory regions. Nature 480: 490–495.

Sun J, Ramos A, Chapman B, Johnnidis JB, Le L, Ho Y-J, Klein A, Hofmann O, Camargo FD. 2014. Clonal dynamics of native haematopoiesis. Nature 514: 322–327.

Thurman RE, Rynes E, Humbert R, Vierstra J, Maurano MT, Haugen E, Sheffield NC, Stergachis AB, Wang H, Vernot B, et al. 2012. The accessible chromatin landscape of the human genome. Nature 489: 75–82.

Wang Y, Xiao M, Chen X, Chen L, Xu Y, Lv L, Wang P, Yang H, Ma S, Lin H, et al. 2015. WT1 recruits TET2 to regulate its target gene expression and suppress leukemia cell proliferation. Mol Cell 57: 662–673.

Wang Y, Yabuuchi A, McKinney-Freeman S, Ducharme DMK, Ray MK, Chawengsaksophak K, Archer TK, Daley GQ. 2008. Cdx gene deficiency compromises embryonic hematopoiesis in the mouse. Proc Natl Acad Sci USA 105: 7756–7761.

Williams K, Christensen J, Pedersen MT, Johansen JV, Cloos PAC, Rappsilber J, Helin K. 2011. TET1 and hydroxymethylcytosine in transcription and DNA methylation fidelity. Nature 473: 343–348.

Wu H, D’Alessio AC, Ito S, Xia K, Wang Z, Cui K, Zhao K, Sun YE, Zhang Y. 2011. Dual functions of Tet1 in transcriptional regulation in mouse embryonic stem cells. Nature 473: 389–393.

Xie M, Lu C, Wang J, McLellan MD, Johnson KJ, Wendl MC, McMichael JF, Schmidt HK, Yellapantula V, Miller CA, et al. 2014. Age-related mutations associated with clonal hematopoietic expansion and malignancies. Nat Med 20: 1472–1478.

Yamazaki J, Jelinek J, Lu Y, Cesaroni M, Madzo J, Neumann F, He R, Taby R, Vasanthakumar A, Macrae T, et al. 2015. TET2 Mutations Affect Non-CpG Island DNA Methylation at Enhancers and Transcription Factor-Binding Sites in Chronic Myelomonocytic Leukemia. Cancer Res 75: 2833–2843.

Yin Y, Morgunova E, Jolma A, Kaasinen E, Sahu B, Khund-Sayeed S, Das PK, Kivioja T, Dave K, Zhong F, et al. 2017. Impact of cytosine methylation on DNA binding specificities of human transcription factors. Science 356: eaaj2239.

Zhang YW, Wang Z, Xie W, Cai Y, Xia L, Easwaran H, Luo J, Yen R-WC, Li Y, Baylin SB. 2017. Acetylation Enhances TET2 Function in Protecting against Abnormal DNA Methylation during Oxidative Stress. Mol Cell 65: 323–335.

Zhu H, Wang G, Qian J. 2016. Transcription factors as readers and effectors of DNA methylation. Nat Rev Genet 17: 551–565.

